# An Implantable Wireless Battery-Free Spatially Selective Vagus Nerve Stimulator

**DOI:** 10.64898/2026.04.10.717669

**Authors:** Edvards Rutkovskis, Enrico Ravagli, Henry Lancashire, Ahmad Shah Idil, Nicole Thompson, Justin Perkins, Ronald Challita, Joseph Hadaya, Umesh Vivekananda, Olujimi A. Ajijola, Kalyanam Shivkumar, Anna Miserocchi, Andy McEvoy, David Holder, Kirill Aristovich

## Abstract

**Objective:** Vagus nerve stimulation (VNS) is an established clinical therapy for drug-resistant epilepsy and other inflammatory conditions. However, off-target stimulation can produce unwanted side effects that limit therapeutic stimulation and hinder the development of new neuromodulation therapies. Selective VNS (sVNS) offers a strategy to reduce off-target organ activation; however, this approach is not available in humans, and no implantable or portable devices exist to trial sVNS in the clinical setup. This work aimed to design, manufacture, and validate an implantable wireless, battery-free stimulator with a selectively addressable output stage for targeted current delivery to specific regions of the human vagus nerve (VN).

**Approach:** We developed a near-field communication-controlled, wirelessly powered, battery-free, temporary implantable multichannel stimulation device, compatible with a 15-channel sVNS cuff electrode (14 selective electrode pairs and one circumferential whole-nerve channel). The device was encapsulated for short-term implantation and evaluated through benchtop characterisation, accelerated ageing, and validation in an acute porcine and a pilot human study.

**Main result:** The sVNS device was evaluated in a porcine (n = 4) trial and a first-in-human pilot study (n = 1). Selective bradycardia of 23.28 ± 12.91% was observed in pigs and 7.5% in the human participant. In the human, a clear separation of bradycardic and tachycardic effects was observed, with additional selectivity in laryngeal activity. Cardiac and laryngeal responses were separated by 231° around the circumference of the nerve.

**Significance:** This work demonstrates the feasibility of wireless battery-free sVNS for cardiac applications using a temporary implantable device. Geometrically selective stimulation has the potential to improve therapeutic efficacy while reducing stimulation-related side effects, and may facilitate future therapies for heart failure and other autonomic disorders.

## 1. Introduction

The vagus nerve (VN) is a key communication link between peripheral organs and the central nervous system (CNS). Approximately 80% of VN fibres are afferent (sensory), while the remaining 20% are efferent (motor). The VN provides extensive innervation to the bronchi, lungs, heart, oesophagus, stomach, liver, and colon, and plays a central role in bidirectional brain–body communication, including the regulation of autonomic function, modulation of inflammatory reflexes, and gut–brain signalling. The VN is comprised of bundles of nerve fibres, called fascicles. Research into the organotopic organisation of the VN focuses on tracing peripheral branches and their associated fascicles [1]. While finer innervation patterns and aspects of the fascicular anatomy have been partially characterised, the fascicular organisation of the VN remains poorly understood, particularly in humans [2–6].

Vagus nerve stimulation (VNS) is a clinically approved therapy for treatment-resistant epilepsy, a condition affecting approximately 20 to 40% of people with epilepsy [2,7]. In addition, VNS is clinically used for treatment-resistant depression. The SetPoint System, an implantable VNS device, has also received FDA approval for the treatment of rheumatoid arthritis [8]. VNS has additionally been investigated as a potential therapy for other inflammatory conditions. More broadly, a range of intractable conditions— such as rheumatoid arthritis, type II diabetes mellitus, and cardiac ischaemia—have been linked to VN activity through diverse mechanisms and have therefore been proposed as potential targets for VNS-based therapies [9–14]. Typically, the therapy involves implantation of a cuff electrode around the left VN and placement of a pulse generator under the clavicle. Conventional VNS therapy stimulates the whole VN; consequently, stimulation amplitude is typically ramped up to the maximum tolerable level, and titrated downwards if side effects become intolerable [15,16]. Activation of the entire nerve results in concurrent stimulation of all organs innervated by the VN rather than of a specific target. Similarly, while transcutaneous electrical nerve stimulation-based approaches provide a non-invasive alternative, they stimulate the nerve non-selectively and therefore inherently lack target specificity. As a result, VNS treatment is associated with multiple side effects, including voice hoarseness, unpleasant skin sensation, shortness of breath or disordered breathing particularly at night, coughing, and headaches [15–18]. The side effects can be mitigated by reducing stimulation current, but at the expense of reduced therapeutic effect. For instance, one possible explanation for the limited efficacy of recent VNS trials for heart failure, such as NECTAR [19], is the wide variability in the stimulation current levels used, which may have prevented delivery of the therapeutic dose required to recruit vagal cardiac efferent fibres. On the other hand, increasing the stimulation intensity of conventional VNS applied to the whole nerve may result in intolerable side effects and/or simultaneous activation of opposing efferent and afferent pathways, resulting in a null effect [20,21].

To address these limitations, various approaches to improve the selectivity of VNS have been developed and are currently under investigation [16]. Some of these include fibre-selective stimulation, spatially selective stimulation, anodal block, kilohertz electrical stimulation, and neural titration. These approaches aim to enhance the precision and therapeutic efficacy of VNS while mitigating stimulation-induced side effects.

Our approach is based on spatially selective VNS (sVNS) and employs a multielectrode cuff architecture comprising 14 equidistant electrode pairs distributed circumferentially around the nerve at the cervical level [22]. sVNS requires identification of the target functional branch, followed by selective stimulation of the corresponding nerve region at the mid-cervical level. Stimulation is applied sequentially through each bipolar channel, defined by one of 14 longitudinally aligned pairs of electrodes, while physiological responses associated with the target are recorded. Once the target region is identified, current is delivered via the independently addressable channel corresponding to the region of interest [1].

Available evidence suggests that cervical VN fascicular organisation is organ-specific [1,23,24]. Consequently, sVNS is expected to improve clinical outcomes and therapeutic efficacy while simultaneously reducing the severity of side effects [22,23,25]. Achieving this requires a targeted stimulation paradigm in which the clinician can select specific nerve sectors and the stimulator can reliably deliver therapy to the corresponding regions. However, to the best of our knowledge, no implantable stimulators have been specifically designed to enable sVNS.

Despite recent achievements in the field of spatial sVNS and the conceptual understanding of its effects [16,22,23,25,26], it is currently implemented using relatively large modular benchtop systems, such as the ScouseTom system, or large current sources connected to switching arrays [27]. Commercial VNS devices for the treatment of epilepsy, such as the Livanova SenTiva, are designed as single-channel stimulators that stimulate the entire nerve. In this work, we present a miniature, 15-channel, wirelessly powered and controlled sVNS device that enables selective stimulation through an implantable platform suitable for short-term preclinical and clinical validation. The acute clinical configuration is sufficiently compact and reliable for deployment in the surgical theatre, enabling evaluation in a real human surgical setting.

One of the major barriers to the development of such systems is the long-term reliability of implanted electronic devices. Any implanted electrical device is exposed to an aggressive physiological environment, including moisture, salinity, and the immune response. As implantation duration increases, so does the risk of device failure due to component degradation or malfunction, often requiring revision surgeries that carry additional clinical risks. Consequently, long-term reliability represents a key safety and design challenge for implantable neuromodulation systems [17]. In the context of VNS, commonly reported technical issues include lead and battery failures, battery depletion, and battery electrolyte leakage, all of which may require device replacement or additional surgical intervention [28,29]. These limitations underscore the need for implantable VNS solutions that prioritise simplicity, robustness, and functional effectiveness. Accordingly, designs that minimise cable length, package volume, and optimise battery characteristics are expected to enhance device longevity and reliability.

Our proposed stimulator design is wireless and battery-free, with the sVNS cuff electrode embedded in the system, allowing implantation of the device near the VN. Since all power is delivered wirelessly through a transmitter, the device can be powered on demand when stimulation is required. A compact acute configuration of the device was developed to enable device validation and perform sVNS in the clinical setting. In this work, we describe the design and manufacturing procedure of the stimulator, followed by a series of benchtop tests to validate and characterise the device. We then present its pre-clinical validation in a porcine model and, finally, a first-in-human (n=1) evaluation.

The aim of this project was to design, manufacture, and test an implantable wireless, battery-free stimulator featuring a selectively addressable output stage, enabling targeted current delivery to specific regions of the VN.

## 2. Methods

### 2.1. Experimental design

We designed a miniaturised sVNS system for use in preclinical porcine experiments and subsequently adapted the system for the first pilot validation of sVNS in humans (Fig. 1).

**Figure 1.**
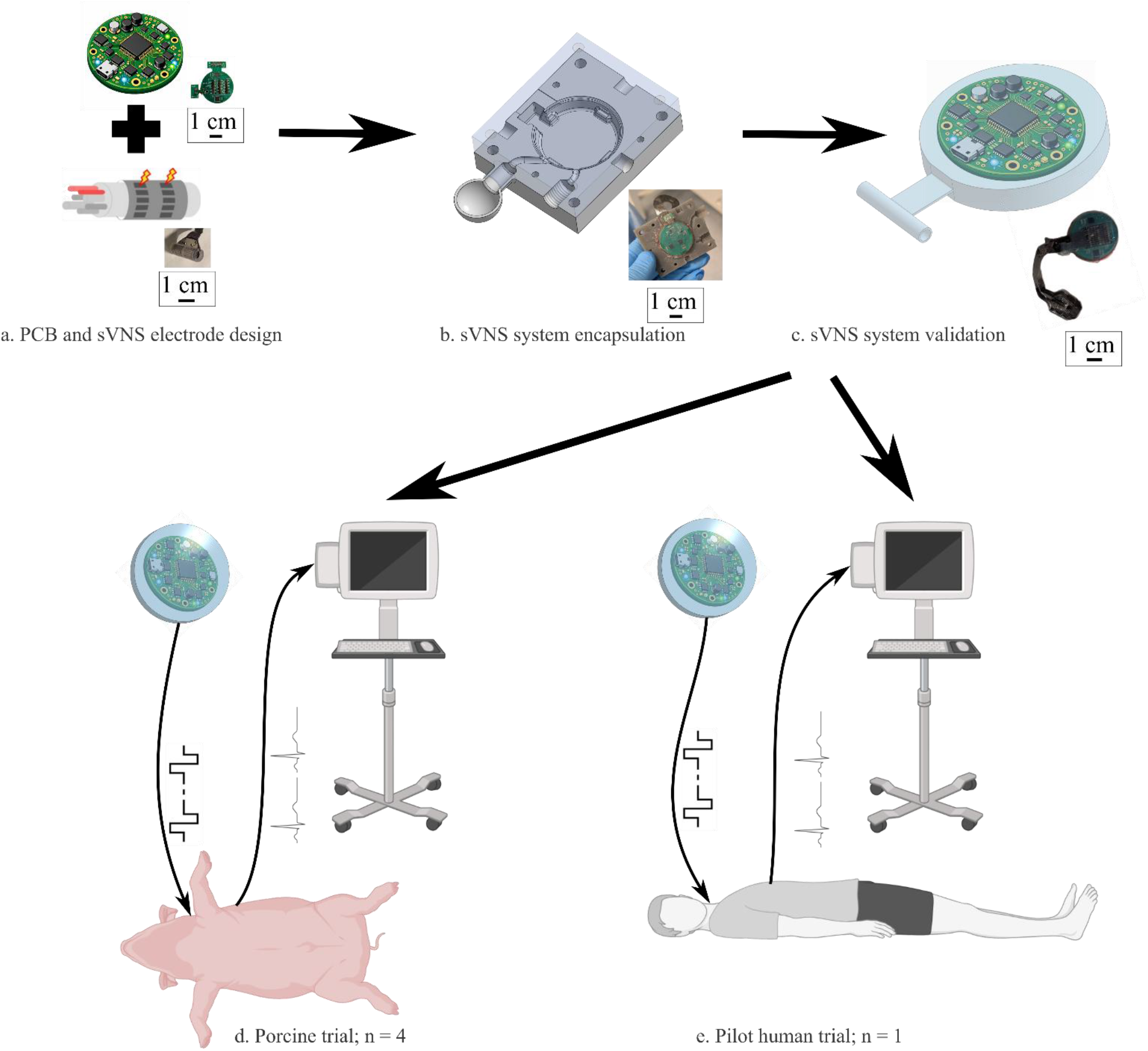
Selective vagus nerve stimulation (sVNS) experimental design. a. A printed circuit board was designed and manufactured using off-the-shelf components. A 14-channel multielectrode cuff array validated in previous works [22,23,30] was employed in the system design. b. The miniaturised stimulation electronics were encapsulated in biocompatible silicone using an injection-moulding technique. c. The system was validated through a series of benchtop tests: the output current was measured across a set of various resistors and in saline, and the biocompatible silicone packaging was assessed in an accelerated ageing test. d. The implantable system was assessed in a porcine trial (n = 4). The sVNS cuff electrode was implanted on either the left or right vagus nerve (VN) in the pigs. A clockwise trial-and-error electrical stimulation paradigm, which utilises longitudinal current steering between two rings of fourteen electrodes, was applied. The protocol comprised 30 seconds of stimulation aimed to recruit cardiac nerve fibres and 30 seconds of stimulation-free window for every 14 channels. Pigs’ heart rate was recorded during the stimulation and stimulation-free time. e. Before the commencement of the selective stimulation in the pilot human trial (n = 1), the entire nerve stimulation was applied to gauge the VN sensitivity to stimulation. Subsequently, the same spatially-selective stimulation protocol was employed. The electromyography activity from the forehead, chin, neck, and shoulder regions was recorded during the stimulation and stimulation-free time windows.

We chose a charge-balanced biphasic square waveform as the primary stimulation waveform based on established evidence demonstrating VN activation with spatial sVNS [1]. The maximum stimulation current was limited to 2 mA to balance safety considerations with sufficient nerve recruitment. A range of pulse width (PW) settings was implemented to enable recruitment of different fibre types. Shorter PW, such as 50 µs in combination with smaller amplitudes, are sufficient for activation of myelinated Aα and Aβ fibres, whereas longer PW in the range of 1 to 4 ms are generally required to activate smaller myelinated and some unmyelinated fibres. The voltage compliance was limited to 20 V to ensure delivery of the programmed current under high contact impedance conditions, which may arise from a small surface area of individual contacts in a multi channel electrode. An additional safety measure was the implementation of wireless power harvesting to eliminate the need for an onboard battery, together with wireless communication to remove any wired interfaces.

The smallest off-the-shelf components were selected during the design stage to miniaturise the overall printed circuit board (PCB) area and device volume, and the system was developed to be fully compatible with a multichannel epineural sVNS cuff electrode array.

The first step of the manufacturing process was PCB production and surface-mounted device (SMD) assembly. After the initial function test, the sVNS multielectrode array (MEA) was soldered to the PCB, and the stimulation output from the cuff electrode was validated in saline. A custom mould was designed and 3D-printed for encapsulating the device in biocompatible silicone. Accelerated ageing tests were performed to test the device’s durability under simulated physiological conditions.

The sVNS system was set to operate in a trial-and-error stimulation mode, stimulating an electrode channel for a programmable time period before switching to the next stimulation channel. The device was implanted at the mid-cervical level in four pigs: on the left side in three animals (n = 3) and on the right side in one animal (n = 1). The device was powered from outside the body using an external near-field communication (NFC) transmitter. Porcine heart rate (HR) was monitored during the stimulation.

The sVNS system was modified as a non-implantable but portable configuration to meet the requirements of the hospital operating theatre environment for use in the human pilot study. Adjusting current amplitude, PW, frequency, and duty cycle is required to ensure recruitment of certain fibre types. For instance, unmyelinated C-fibres require higher currents and longer PWs than myelinated A-fibres. Therefore, prior to the selective stimulation paradigm in the human participant, the whole VN was stimulated using a pair of fully circumferential electrodes located in the same cuff to confirm the nerve sensitivity to the stimulation current. The aforementioned trial-and-error stimulation was subsequently performed. Human vital signs, including the HR and surface electromyography (EMG) activity, were recorded.

### 2.2. sVNS system specification and characterisation

The device was designed to deliver a maximum of 2 mA biphasic current as a stimulation current source to evoke a physiological response through modulating the VN activity. The desired current source voltage compliance was estimated based on stainless steel electrode conductivity parameters [22]. The worst-case voltage compliance was estimated at 20 V for platinum-coated stainless-steel electrodes, for which the electrode-tissue impedance can reach values of up to 10 kΩ. Based on these considerations and prior studies, the technical requirements (Table 1) were defined [16,22].

**Table 1.** sVNS stimulator technical specification.

| Parameter | Value |
| --- | --- |
| Waveform | Biphasic Pulse |
| Amplitude | 2.026 mA (in steps of 32 μA) |
| Pulse width | 50 μs, 1 ms, 2 ms, 4 ms |
| Pulse frequency | 1, 20, 50 Hz |
| Number of stimulation channels | 14 selective stimulation channels + 1 entire nerve stimulation channel |
| Power and communication | Wireless – NFC |
| Size | $\varnothing 33.73 \text{ mm} \pm 0.15 \text{ mm}$ |

The sVNS system current injection performance was measured by setting different current levels via the NFC interface and measuring the voltage over 100 Ω in series with the corresponding channel. Voltage compliance was confirmed by changing saline concentration and measuring voltage across the channel (Fig. 4c). Five measurements were performed for each representative current setting (0.515, 0.965, 1.447, and 2.026 mA), pulse width (50, 150, 1000, 2000, 4000 µs) and pulse frequency (1, 5, 10, 20, 50 Hz). The mean value, standard deviation, percentage error with respect to the programmed value, and mean percentage error were calculated.

The injected charge was calculated as the cumulative sum of the current injected into the saline, multiplied by the sampling period, as shown in equation (1). The calculation was performed for all tested saline concentration levels (Fig. 4d). The background noise accumulation was subtracted from the sum in each case to compensate for the background offset.

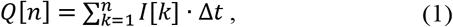

where *Q*[*n*] is the injected charge at sample *n, I*[*k*] is the measured current at sample *k*, and Δ*t* is the sampling period (250 ns).

The current consumption of the sVNS system was measured by powering the system with a CR2450 3 V lithium coin cell battery and recording the current drawn from the battery.

### 2.3. sVNS system design

#### 2.3.1. System architecture

The sVNS system (PCB diameter 33.73 mm ± 0.15 mm, n = 3) uses an NFC interface to communicate with an external programming device (PN532, Adafruit) and receive operational and stimulation parameters. The system harvests all required power from the same NFC interface via a 13.56 MHz-tuned antenna. The device architecture (Fig. 2) is based on discrete off-the-shelf components. These include an NFC integrated circuit (IC) (NT3H2211, NXP Semiconductors, Netherlands), responsible for communications and power harvesting, a voltage regulator (LT8410, Analog Devices, USA) that boosts harvested voltage to 20 V and supplies the high-voltage circuitry, an operational amplifier (OPA990, Texas Instruments, USA), a 20 V analogue switch to provide charge-balanced current switching (MUX36D08, Texas Instruments, USA), and two 20 V analogue switches for output waveform demultiplexing (MUX36S16, Texas Instruments, USA). Silicon Lab’s 8-bit EFM8 Sleepy Bee was chosen as the main microcontroller unit (MCU) due to the available package size, current reference function, and low power requirement; the system’s stimulation architecture was based on a modified ReStore nerve stimulation system [31]. The stimulator firmware was developed in C and implemented using the Simplicity Studio 5 integrated development environment (Silicon Labs, USA).

**Figure 2.**
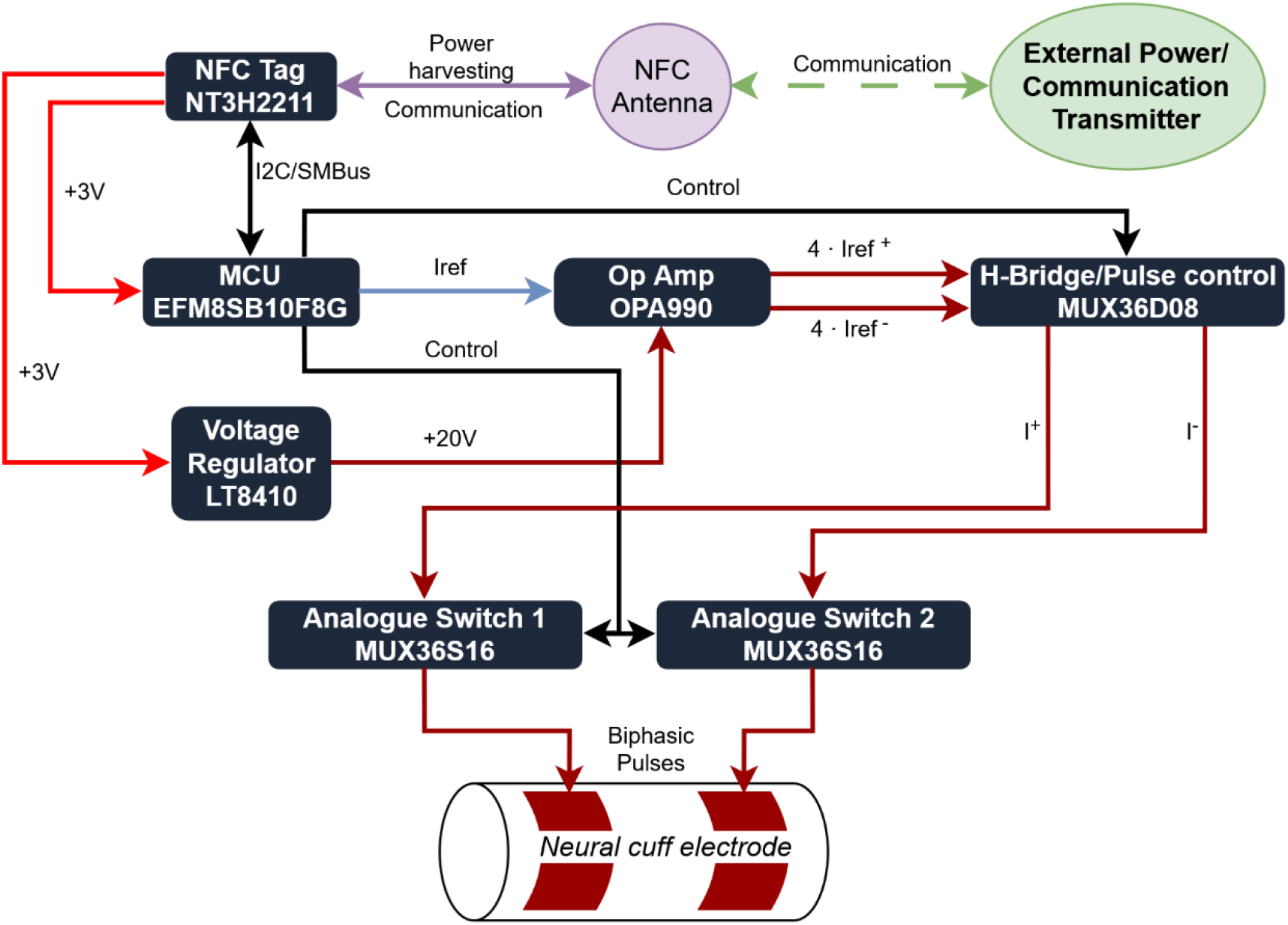
Selective vagus nerve stimulation system architecture. An external near-field communication (NFC) transmitter is utilised to wirelessly communicate and power the stimulator. The harvested radiofrequency energy and programming data are received by an NFC integrated circuit via an NFC antenna and communicated to the microcontroller through an I^2^C (Inter-Integrated Circuit) interface. The microcontroller configures the stimulation parameters, including current amplitude, pulse polarity via an analogue H-bridge, and channel selection through output analogue switches. A boost regulator converts the harvested voltage to 20 V to enable high-voltage compliance for the multichannel current-controlled current source. The boosted supply also powers the switching network to enable high-voltage multichannel stimulation.

The PCB was designed using EAGLE (Autodesk, USA). The four-layer circular PCB fabrication and component assembly were performed by PCBWay (China).

#### 2.3.2. sVNS system multichannel stimulation array

The multichannel array (Fig. 3a) comprised 14 independent stimulation channels, each formed by a pair of coaxial, longitudinally placed electrodes, each sized 3.00 × 0.35 mm^2^ (Fig. 3d, 3e). Additionally, a pair of full circumferential VN stimulation electrodes, each sized 9.18 × 0.5 mm^2^, was incorporated in the MEA and used to perform stimulation of the entire VN. The cuff array consisted of platinum-coated stainless-steel foil embedded in biocompatible silicone. The electrode longitudinal position, shape, and manufacturing technique are informed by previous works by our group [22,32]. The longitudinal stimulation activates fibres that are adjacent to the stimulation electrodes, thus spatially selecting VN fibres and fascicles over the cross-section of the nerve.

**Figure 3.**
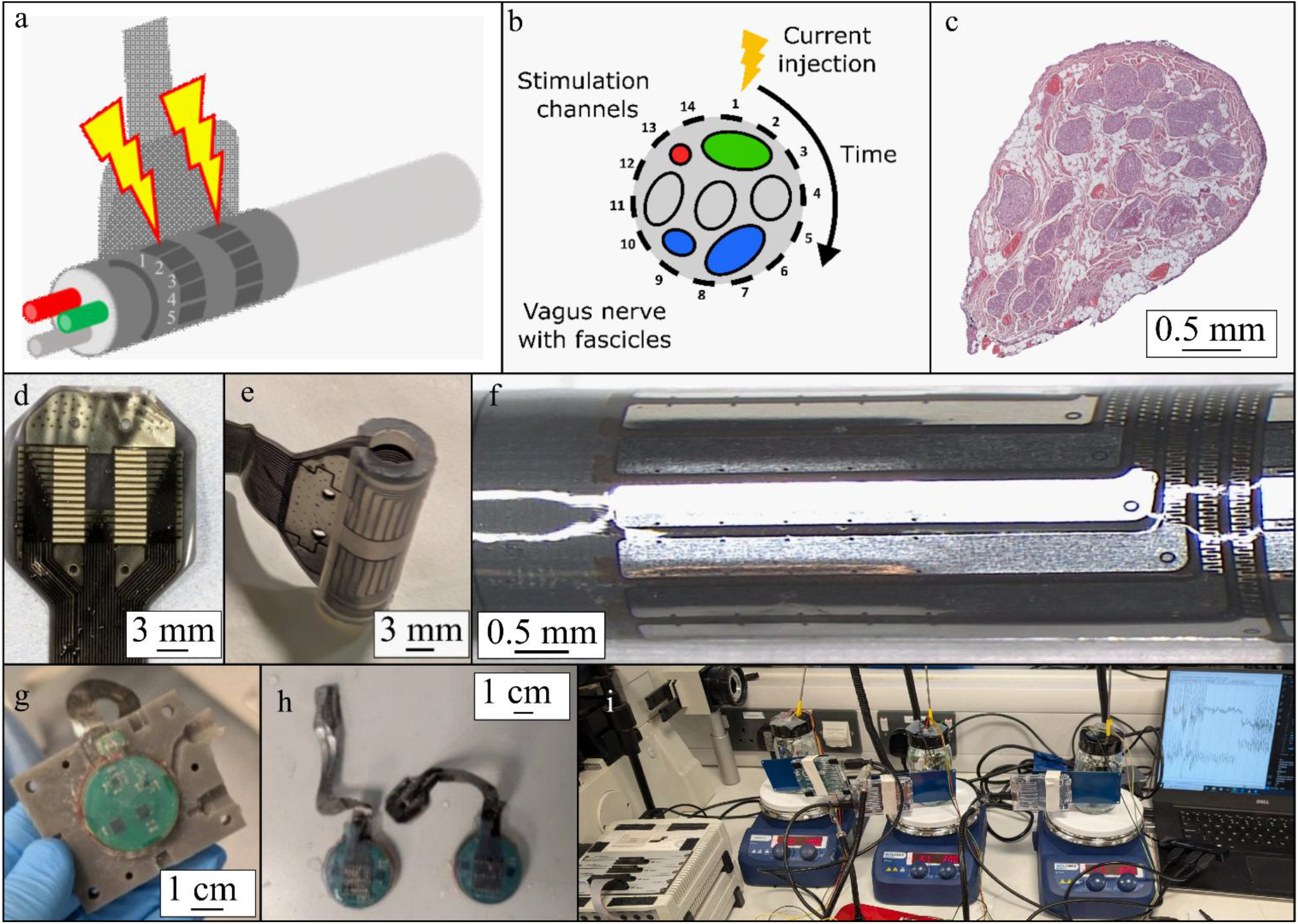
Overview of the selective vagus nerve stimulation (sVNS) protocol, device design, nerve cuff electrode fabrication, and device characterisation in the accelerated ageing setup. a. A schematic of sVNS cuff electrode, placed on a vagus nerve (VN). The cuff is shown delivering stimulation current between the first pair of longitudinally placed electrodes. b. VN cross-section shows organ-specific fascicles; each colour represents separate organ innervation. Numbers from 1 to 14 depict channels within the stimulation cuff wrapped around the VN. The stimulation protocol encompasses a longitudinal stimulation of a single channel at a time. c. Cross-section of the porcine VN showing individual fascicles—small nerve bundles that exhibit organ-specific organisation in pigs and are thought to have a similar organisation in humans [23]. d, e. An unwrapped sVNS stimulation cuff electrode and the cuff electrode with the distal stimulation side encapsulated in a cylindrical configuration. f. Close-up view of a coated electrode side, showing the patterned conductive track that enables mechanical extensibility. g. A 3D-printed injection mould with the encapsulated sVNS device. h. Encapsulated sVNS devices. i. Accelerated ageing setup utilised in the assessment of the silicone encapsulation.

The manufacturing procedure was as follows. First, a glass slide was covered with biocompatible silicone (MED-4220, NuSil Technology) and cured. A stainless-steel foil was then rolled on top of the silicone layer. The array shape was formed by cutting the outline with a laser. The electrode tracks were further patterned to improve the cuff’s stretchability and compliance with respect to the nerve tissue. Next, the second layer of silicone was applied on top of the array and laser-cut once more. The electrode pads were patterned to improve the contact impedance and the platinisation layer adherence. The excess silicone was then peeled off. The electrodes were platinised by electrodeposition. The electrode’s proximal end was connected to the custom female 30-pin header or soldered directly to the 30-pin male header of the sVNS device. To remove residual platinum solution and contaminants, the cuff electrode was cleaned in an ultrasonic bath for 5 minutes in isopropyl alcohol, followed by 5 minutes in deionised water. After manufacture, the distal end of the flat array (Fig. 3e) was embedded in a custom-made biocompatible silicone tube to give it a cylindrical shape (Fig. 3f). After embedding within the silicone tube, the cuff was further cleaned by sonication in deionised water for 5 minutes and rinsed with deionised water.

Prior to the human trial, the cuff electrode was sterilised in the surgical-grade sterilisation department at the 121°C sterilisation cycle.

The cuff electrode is designed to be placed on the cervical section of the VN (Fig. 6a) by opening the cuff with the handling pad, sliding the distal side of the cuff under the nerve, and releasing the handling pad.

#### 2.3.3. Communication and power harvesting antenna

Communication and power harvesting were implemented using an NFC integrated circuit (NT3H2211) operating at 13.56 MHz. The antenna connected to the IC antenna pins was tuned with parallel capacitors to resonate at 13.56 MHz. Resonance was measured using the impedance magnitude peak and phase transition using an impedance analyser (6500B, Wayne Kerr Electronics) [33]. A 0.5 mm thick, single-turn, enamelled copper wire was used as the antenna coil. The coil was tightly wrapped around the PCB and tuned to resonance at 13.56 MHz using two surface-mount capacitors (nominal 120 pF and 560 pF) connected in parallel. Series resistance and parasitic reactance were minimised with short antenna leads. After successful tuning, defined as the characteristic peak in impedance magnitude around 13-14 MHz, the antenna was soldered to the antenna pins, and communication with the sVNS system was verified using the transmitter. Further tuning was achieved by selecting appropriate alternative parallel capacitor values.

#### 2.3.4. sVNS system encapsulation

Before encapsulation, the multichannel stimulation array was soldered directly to the 1.27 mm 30-pin sVNS output connector (three 10-pin M50-3631042 connectors, Harwin). The communication and power harvesting antenna was soldered to the antenna input pads, forming a single integrated system unit ready for silicone encapsulation.

A custom 3D-printed mould was designed to encapsulate the system in a biocompatible package (Fig. 3g). The encapsulation procedure was performed in an ISO 6 standard cleanroom.

The sVNS system PCB was submerged and washed in an ultrasound cleaner for 5 minutes in acetone, 5 minutes in isopropyl alcohol, 5 minutes in deionised water, and finally rinsed in deionised water for 2 minutes. The system was dried using nitrogen gas and further dried in an oven at 150°C for 15 minutes, or until no visible moisture remained.

An optically clear two-part silicone elastomer (MED-4220, NuSil Technology) was used to encapsulate the system using a custom 3D-printed high-temperature resin mould. The mould was cleaned using the same procedure as the VNS system PCB. Equal masses of parts A and B (5 g each) were combined to prepare the silicone mixture. The components were mixed in a vortex mixer (SpeedMixer DAC FVZ-K, Hauschild) for 3 min at 2700 RPM and subsequently injected into the mould at a pressure of 2 bar. The mould was centrifuged in a custom-built vacuum centrifuge at 700 RPM and 25 mbar pressure to remove trapped air and ensure conformal silicone coating, including voids beneath soldered surface-mount components. Finally, the encapsulated device was cured in an oven at 150 °C for 40 min and checked for complete curing and absence of voids. The encapsulated device (Fig. 3h) was rinsed in IPA and deionised water for 2 minutes.

### 2.4. Accelerated ageing model and setup

Accelerated ageing tests were conducted to evaluate the performance of the encapsulated device under conditions simulating long-term implantation in vivo. The acceleration factor given in equation (2) was derived from the Arrhenius equation and used to estimate the equivalent ageing duration under elevated temperature conditions [34].

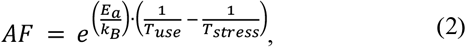

where *AF* is the acceleration factor, *E*_*a*_ is the activation energy of the dominant ageing mechanism, *k*_*b*_ is the Boltzmann constant (8.617 × 10^−5^ eV K^-1^), *T*_*use*_ is the nominal use temperature (37 °C, 310 K), and *T*_*stress*_ is the elevated stress temperature (67 °C, 340 K). As the activation energy of the dominant degradation mechanism of silicone encapsulation is not precisely established, a conservative value of *E*_*a*_ = 0.7 eV was selected based on prior implantable polymer ageing studies [34,35]. The stress temperature of 67°C was chosen to represent an acceleration scenario for this activation energy [35]. The model with these parameters yields the accelerated ageing factor *AF* = 10.1, implying that each day of the experiment would approximate 10 days *in vivo*. We note this acceleration factor because: *E*_*a*_ is unknown and small errors in *E*_*a*_ cause large errors in *AF*; the Arrhenius model depends on the assumption that device failure is attributable to the same failure mechanisms at *T*_*use*_ and *T*_*stress*_; and the failure mechanism may be one to which the Arrhenius model is poorly suited, such as mechanical failure modes [35,36].

The accelerated ageing setup (Fig. 3i) comprised a glass tank filled with phosphate-buffered saline, a heater-stirrer (SCI280-Pro, Scilogex), and the sVNS device immersed within the glass tank and wirelessly powered by an external transmitter (PN532, Adafruit) positioned outside the tank. The tank incorporated small apertures for the temperature feedback probe and recording electrodes to capture the stimulation output during the experiment. The hotplate with built-in magnetic stirrer was programmed to maintain 67°C with a stirring speed of 200 RPM.

sVNS devices (n = 3) were tested continuously until failure for up to 15 days. Device memory readouts, containing programming parameters and the current stimulation channel, were performed daily. Additionally, stimulation waveforms were recorded during the experiment to determine the time of failure. The devices were set to continually stimulate at the highest available stimulation settings (I = 2026 µA, PW = 4000 µs, PF = 20 Hz, 30 seconds on and 30 seconds off continuous working mode). The test commenced once both the temperature and stirring setpoints were achieved.

### 2.5. Device validation in an in vivo porcine model

A clockwise trial-and-error electrical stimulation paradigm (Fig. 3b) was used in the porcine trial. The multichannel electrode array circumferentially positioned around the VN enabled stimulation of discrete nerve sectors, thereby recruiting specific fascicular regions (Fig. 3c) rather than the entire nerve. When the sVNS system stimulated a channel adjacent to the fascicles of interest, a corresponding fascicle-dependent physiological response was observed on the physiological recording system.

The sVNS system was tested on anaesthetised pigs at UCLA (male Yorkshire, S&S Farms Roman, CA, USA, 53.7 ± 3.7 kg, n = 4) to study the physiological effects of selective stimulation. The implantable device porcine trial was conducted in the same animals used in the spatially selective cardiac efferent neuromodulation study at UCLA previously reported in [1]. The porcine study was conducted in accordance with the guidelines set by the University of California Institutional Animal Care and Use Committee and the National Institutes of Health Guide for Care and Use of Laboratory Animals. All experiments were approved by the UCLA Chancellor’s Animal Research Committee (approval no. ARC 2016-085). The animals were housed in the vivarium with ad libitum access to water and were fed twice daily by vivarium staff.

The same surgical, anaesthesia, and physiological monitoring protocols were utilised as described in our previous work [1]. Tiletamine‐Zolazepam (4 to 6 mg/kg) was used intramuscularly to induce general anaesthesia in all four animals. The anaesthesia was maintained with vaporised isoflurane, intubated with an endotracheal tube. A continuous intramuscular injection of fentanyl (0.2 μg/kg/min) was administered to maintain analgesia for the duration of the whole experiment. The animal was ventilated using pressure control mode for the duration of the implantable sVNS trial. The surgical region was aseptically prepared, and a 5 cm to 7 cm region of the left VN and right VN was exposed using a blunt dissection technique. sVNS device implantation side was selected based on the timeslot availability of the main experiment [1]. The selective stimulation cuff electrode was placed on the pig’s left or right VN (Fig. 6a) by opening the access slit and sliding the VN inside the cuff. The pulse generator was positioned subcutaneously within the surgical pocket. The left VN was selected as the implantation target in pigs 1, 2, and 3. In pig 4, the right VN was targeted instead. Following electrode implantation, anaesthesia was transitioned to alpha-chloralose to minimise depression of cardiac and autonomic reflexes (initial bolus of 50 mg/kg, followed by 25 to 50 mg/kg/h, i.v.). Vital parameters were measured during the experiment, and invasive blood pressure was recorded using an ActiCHamp Plus amplifier (Brain Products, Germany) with a sampling rate of F_s_ = 50 kHz. Data were further processed using MATLAB (MathWorks, Natick, MA, USA).

During experiments, the sVNS system stimulated the pigs’ VN using biphasic current pulses. A pulse width of 1 ms, pulse frequency of 20 Hz, and amplitude of 1 mA were used for the cardiac sVNS protocol to elicit a bradycardic response through the activation of efferent cardiac fibres. Channel stimulation and subsequent pulse-free time were 30 seconds, resulting in channel switching every 60 seconds.

### 2.6. Pilot human trial

A modified sVNS system was employed (Fig. 6c) in a pilot sVNS human trial. An acute clinical configuration was developed to address the safety, portability, and sterility required by the surgical setup. The sVNS device was fitted in a custom 3D-printed case with the transmitter embedded in the case. The transmitter was connected to a 5 V USB port of a laptop, containing a custom graphical user interface developed in Python. The sVNS device was powered wirelessly, achieving full separation from the mains voltage. The sVNS system generated 5V TTL trigger signals through a D-Sub interface to the EMG amplifier to mark the stimulation onset and offset.

Ethical approval for this clinical trial was granted by the NHS Research Ethics Committee (REC reference number 22/LO/0463). The trial is registered with the Integrated Research Application System (IRAS) under project ID 276275 and is publicly accessible via ClinicalTrials.gov (identifier NCT05664854).

A candidate for VNS therapy, scheduled to receive a LivaNova SenTiva VNS system (comprising both the IPG and the stimulation electrode), was consented and recruited for the trial. The following describes the general anaesthesia protocol. Anaesthesia was induced using a combination of 1% propofol (200 mg) and sevoflurane. The patient was positioned supine. A neuromuscular blocking agent (rocuronium) was administered by the surgical team immediately after the onset of the anaesthesia, and its effects were reversed using sugammadex (200 mg) 6 minutes prior to the start of the sVNS trial.

Five surface EMG electrodes (Neuroline Ground, Ambu, Malaysia) were placed on the patient at the forehead, chin, mid-neck, and bilaterally over the shoulders, and were connected to the ActiCHamp Plus recording headbox. Following electrode placement, non-surgical areas were covered with antimicrobial surgical drapes by the theatre staff. An additional surgical drape was placed over the supporting metal table with the clinical sVNS system.

Once the anaesthesia was successfully induced and maintained, and the EMG electrodes were placed, the surgeon identified the surgical site using anatomical landmarks. The temporary sVNS cuff electrode was placed around the left VN by opening the silicone tube support, wrapping the cuff around the VN, and then releasing the silicone tube. The cuff opening was placed ventrally. Anatomically, the cuff position is ∼5 cm distal to the nodose ganglion and ∼5 cm proximal to the clavicle. After the sVNS cuff electrode was placed, a 30-minute experiment window commenced. A short stimulation of the entire VN was delivered, with the current amplitude increased incrementally from the starting value of 0.5 mA until a change in HR (tachycardia or bradycardia) was observed. For this patient, an HR change was observed at 1.6 mA; this current amplitude was therefore selected for the cardiac stimulation protocol. The previously described clockwise trial-and-error spatially selective VNS (Fig. 3b) of the left VN was then performed at 1.6 mA.

To ensure separation between the sVNS system and the system operator, as well as the surgically sterile zone during VNS surgery, a sterilizable extension cable connecting the sVNS system to the cuff electrode was developed and utilised during the surgery.

### 2.7. Data recordings and analysis

An ActiCHamp Plus high-specification EEG amplifier (Brain Products, Germany) was utilised to capture physiological data with a sample rate of 50 kHz.

HR percentage change was calculated using HR derived from invasive left ventricular pressure recordings sampled at 50 kHz in the porcine experiments, and from the surface EMG recordings in the pilot human study. For the former, the third-order Butterworth 40 Hz low-pass filter was applied to isolate QRS complexes. A zero-cross algorithm was applied, and the interbeat distance was calculated and converted to the instantaneous HR in beats per minute. For the latter, R-wave detection was applied, and the distance between the detected R waves was measured and converted to the instantaneous HR in beats per minute.

Since HR is inherently defined only at beat occurrences, instantaneous HR values were first associated with the timing of the corresponding R-peaks. To obtain a continuous heart-rate time series aligned with the original sampling rate, the instantaneous HR values were resampled using a zero-order hold (sample-and-hold) interpolation, whereby each HR value was held constant until the next detected beat. The average HR of the stimulation window and the subsequent no-stimulation control window were calculated. An HR percentage change was calculated as the ratio of the average HR within the stimulation window and the stimulation-free control window (equation 3). The mean HR percentage change for each channel was calculated across animals (n = 4) and is presented as mean ± standard error of the mean (SEM).

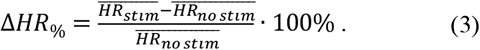

The implantable system was compared to the benchtop system (ScouseTom [27]) used in our previous sVNS study [1] by applying both systems to the same animals and calculating the resulting differences in HR evoked by stimulation of the corresponding VN. A paired-sample t-test was applied to the mean HR change evoked by the stimulation of corresponding channels in both systems to determine whether the difference between the two systems was statistically significant at α = 0.05.

To calculate the averaged evoked potential (EP) value for the surface EMG recording in the pilot human trial, the epoch of the EMG activity within 50 ms of the sVNS biphasic stimulation was averaged for the whole stimulation period (5 seconds for the laryngeal activation protocol with PW of 50 µs, 30 seconds for the cardiac protocol with PW of 1 ms). The EP was defined as the characteristic peak-to-peak EMG activity within 5 to 20 ms after the stimulation of the VN. The EP is attributed to the sVNS-induced laryngeal motor EP [37]. All detected peak-to-peak values of EPs were arithmetically averaged to calculate a single representative EP average per stimulation window. The chin electrode was used for this calculation because it enabled more reliable automated EP peak detection compared to the neck electrode.

## 3. Results

### 3.1. sVNS system functional tests in saline

The multichannel sVNS device showed consistent performance both on a bench resistor test and with the stimulating electrode array submerged in saline (Fig. 4a – d). Representative parameters were programmed, and the output, with respect to maximum stimulation current, minimal biphasic PW, and pulse frequency, matched the desired specification (Table 2). The largest observed current deviation was ±0.006 mA for the 0.965 mA setting, with the highest error of 3.83% for the same setting.

**Figure 4.**
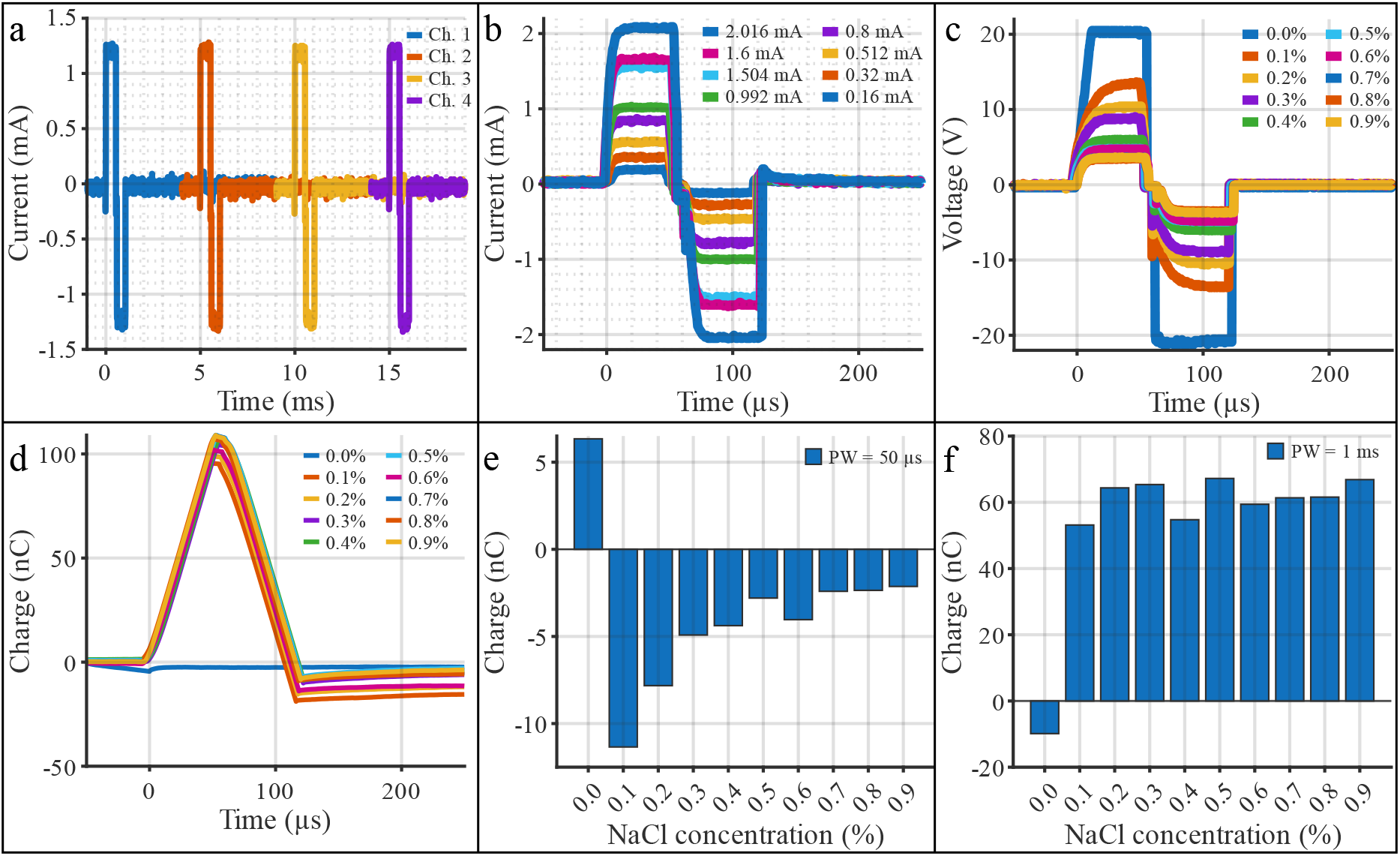
Selective vagus nerve stimulation (sVNS) system functional test results. a. Multichannel current injection into 0.9% saline using the sVNS cuff. Channels switched every 5 ms, biphasic pulse width (PW) was 0.5 ms, programmed current amplitude was 1.2 mA. b. Variable current level tests in 0.9% saline with a 14-channel sVNS cuff. Biphasic current injection was performed in the first channel (PW = 50 µs). c. Measured voltage excursion of the 14-channel selective cuff electrode at various NaCl concentrations (PW = 50 µs). d. The injected charge was calculated from the injected current in different saline concentrations. Charge imbalance after the negative phase can be observed. e, f. Net injected charge at 1 mA current amplitude after 50 µs biphasic pulse and 1 ms biphasic pulse, respectively.

**Table 2.** Programmed and measured stimulation parameters.

| Parameter | Programmed value | Measured mean (±S.D.)<sup>1</sup> | Percentage error (%) |
| --- | --- | --- | --- |
| Stimulation current amplitude (mA) | 0.515 | 0.525 (±0.004) | 1.94 |
| | 0.965 | 1.002 (±0.006) | 3.83 |
| | 1.447 | 1.474 (±0.002) | 1.87 |
| | 2.026 | 1.975 (±0.004) | 2.52 |
| Biphasic PW (μs) | 50 | 55.4 (±0.4) | 10.88 |
| | 1000 | 1069.2 (±5.4) | 6.92 |
| | 4000 | 4272.0 (±30.3) | 6.80 |
| Frequency (Hz) | 1 | 0.9 (±0.002) | 6.04 |
| | 20 | 20.6 (±0.000) | 1.26<sup>2</sup> |
| | 50 | 47.3 (±0.170) | 5.37 |

<sup>1</sup>Current injected using an sVNS cuff (channel 1) submerged in 0.9% saline, PW 50 μs, PF 20 Hz.
<sup>2</sup>The firmware was specifically optimised for 20 Hz stimulation in the clinical trial.

The stimulation switching function conformed to the desired parameters, allowing seamless switching between channels in less than 100 µs, allowing singular pulses to be delivered to the required channels (Fig. 4a).

Performance of various current settings is shown (Fig. 4b) to demonstrate the stability of current delivery within the minimum programmable PW (50 µs). PW setting precision and consistency were satisfactory, with the mean error of 10.88% and 6.92% for the 50 µs and 1 ms settings.

To ensure charge-balanced current delivery to the tissue, electrodes were shunted prior to the positive phase, after the positive phase, and before the negative phase, and after the negative phase until the delivery of the next pulse. This approach prevents accumulation of charge and electrode polarisation during or after any stage of the injection. Despite this, a charge imbalance after negative phase injection was observed (Fig. 4d). This is attributed to charge accumulation on the 10 µF DC-blocking capacitor and its subsequent discharge after the electrode shunting phase. For 0.9% NaCl, the typical net injected charge after a biphasic pulse was −2.1 nC for a 50 µs, 1 mA waveform and 68 nC for a 1 ms, 1 mA waveform (Fig. 4e,f).

These correspond to 1.86% and 3.13% of the total charge delivered per biphasic pulse, respectively. The system was powered wirelessly through water up to a coil-to-coil distance of 10 mm. Calculated mean implant power consumption was 11.946 mW (Table 3).

**Table 3.** sVNS system current consumption.

| Parameter | Mean (± S.D.) |
| --- | --- |
| Idle current (mA) | 2.229 (±0.052) |
| Mean current<sup>1</sup> (mA) | 3.982 (±0.052) |
| Peak stimulation current (mA) | 5.735 (±0.052) |

<sup>1</sup>Calculated as $I_{mean} = I_{idle} \cdot D + I_{peak} \cdot (1 - D)$ , where D is the duty cycle.

The total cost of PCB manufacture and component assembly for a single device was USD 97.18. This cost excludes sVNS array production and device encapsulation.

### 3.2. Accelerated ageing results

To test the device encapsulation, sVNS devices (n = 3) were subjected to accelerated ageing in phosphate-buffered saline at 67°C while stimulating at the maximum current amplitude and PW at 20 Hz. The first failure of an sVNS device occurred after 10 days in the accelerated ageing experiment (Table 4). The fault condition was identified by the loss of communication with the sVNS system, accompanied by cessation of the stimulation. Visible traces of corrosion were detected near the antenna terminals after the device’s post-experiment inspection. The second device failed after 11 days. The third device survived for the duration of the experiment (15 days).

**Table 4.** Accelerated ageing result summary.

| Day | Time | Test result | Notes |
| --- | --- | --- | --- |
| Day 1 | 16:00 | Device 1,2,3: Pass | Start of the experiment |
| Day 10 | 16:00 | Device 1: Fail<br>Device 2, 3: Pass | T=67°C, 200 RPM |
| Day 11 | 18:00 | Device 1,2: Fail<br>Device 3: Pass | T=67°C, 200 RPM |
| Day 15 | 18:00 | Device 1,2: Fail<br>Device 3: Pass | End of the experiment |

If our estimated accelerated ageing factor of *AF* = 10.1 holds, 10 days in the accelerated ageing setup corresponds to a theoretically predicted 100 days *in vivo*. As previously noted, this prediction relies on assumptions which may not hold true, including: the activation energy, *E*_*a*_; that device failure mechanisms are the same at *T*_*use*_ and *T*_*stress*_; and that the Arrhenius model is applicable to the device failure modes. We do not claim a specific device lifetime at 37°C service temperature.

### 3.3. Porcine in vivo experiment results

Four porcine experiments were performed using the sVNS system to evaluate its cardiac effects. A single animal HR response to stimulation of the right VN is shown (Fig. 5a). In this experiment, the stimulation parameters were 1 mA, PW of 1 ms, and pulse frequency of 20 Hz, which targeted activation of cardiac efferent fibres. A 58% HR decrease, corresponding to an HR drop from 104 BPM to 13 BPM, was achieved in channel 1.

**Figure 5.**
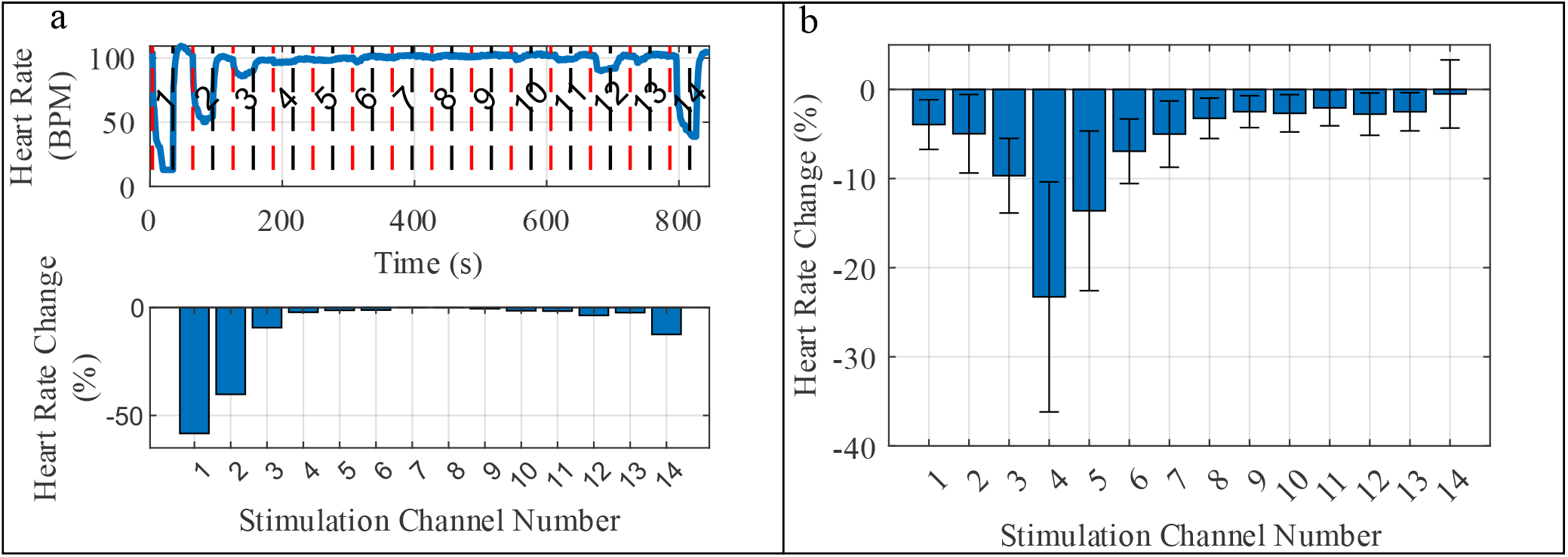
a. Representative raw heart rate (HR; top) and corresponding calculated HR percentage change (bottom) of one of the pigs during the cardiac fascicle stimulation. A numbered dashed red line signifies the onset of the stimulation period, and the unnumbered black dashed line signifies the cessation of the stimulation-free period. b. Mean HR percentage change ± standard error of the mean across all porcine trials (n = 4), after aligning the maximal HR decrease to channel 4.

**Figure 6.**
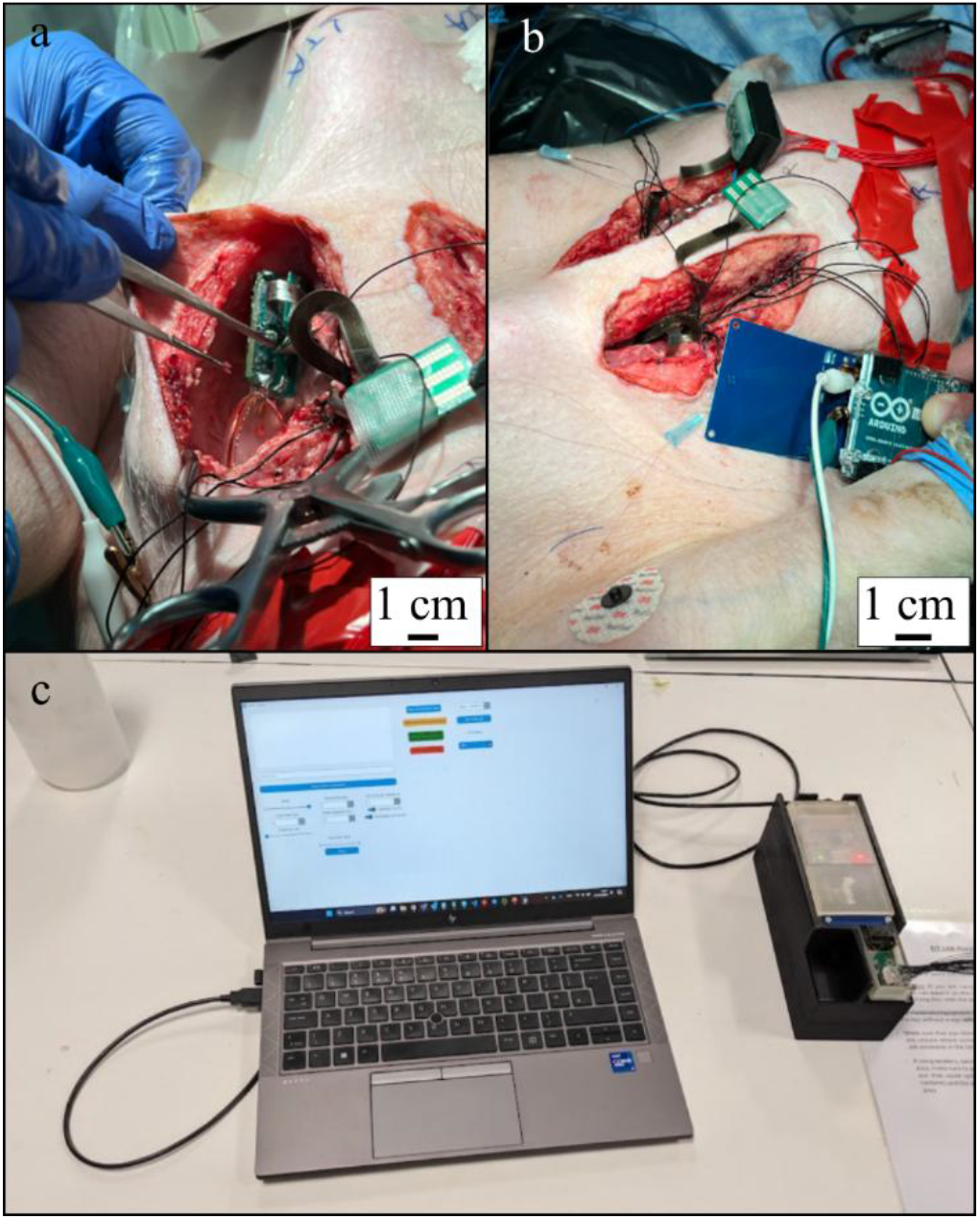
Selective vagus nerve stimulation (sVNS) device applications in the *in vivo* studies. a. sVNS system implantation in a pig’s cervical vagus nerve section. b. An implanted sVNS system powered by an external near-field communication (NFC) transmitter, demonstrating wireless communication and power transfer to the implant. c. Experimental clinical sVNS system, based on the implantable sVNS device, for the pilot human trial. The system includes the sVNS pulse generator (square printed circuit board form factor), NFC communication and power system, a sterilisable cuff electrode extension, and a laptop with a graphical user interface.

Analysis of the mean HR decrease across animals was performed after rotationally aligning all stimulation channel data according to the largest stimulation response of the first animal, which was observed in channel 4. Mean values ± SEM are reported. A selective HR decrease of 23.28 ± 12.91% was achieved in all animals (n = 4; Fig. 5b) while stimulating the left VN with 1 mA current and 1 ms PW at 20 Hz, and the right VN with 2 mA, 1 ms PW at 20 Hz. Individual HR responses are presented in the Supplementary Material (Fig. S1 and S2). The paired-sample t-test did not reveal any statistically significant differences in stimulation-induced HR change between the benchtop and the implantable system across all channels (Supplementary Table S1). The smallestp-value was observed for channel 7 (p = 0.214), whereas the largest p-value was observed for channel 1 (p = 0.901).

### 3.4 Pilot human trial result

No adverse effects related to sVNS were reported following the surgery and postoperative recovery period. Processed HR data indicate a decrease in HR during stimulation of specific channels, but not others (Fig. 7a). In particular, stimulation of channels 8 to 10 resulted in a reduction of HR, with the most pronounced decrease (7.5%) observed at channel 10. In contrast, stimulation of other channels (1 to 7 and 11 to 14) resulted in a mild HR increase (0.92% to 3.83%). Throughout stimulation, the heart remained in sinus rhythm. We speculate that the distribution and pattern of the observed cardiac effects suggest that cardiac efferent fibres are localised in proximity to channels 8 to 10, whereas cardiac afferent fibres are distributed near channels 1 to 7 and 11 to 14.

**Figure 7.**
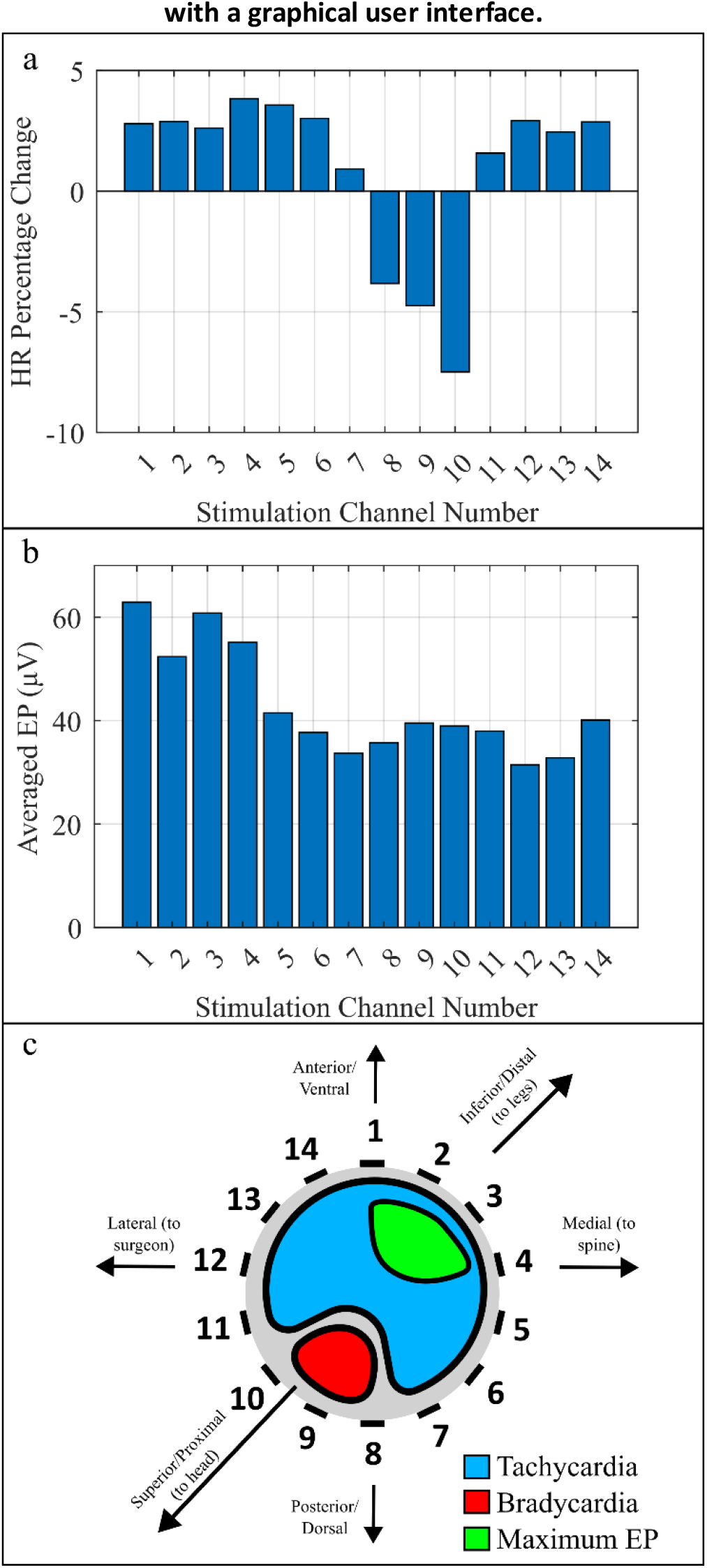
Selective vagus nerve stimulation (sVNS) shows selective stimulation resulting in either tachycardia or bradycardia in a human participant. a. sVNS system used in a successful pilot human trial, showcasing a heart rate (HR) decrease of 7.5% (channels 8-10) as opposed to a minor 2-4% HR increase (channels 1-7 and 11-14). b. Averaged laryngeal activity during the cardiac stimulation protocol measured from the electromyogram, showing the highest evoked potentials in channels 1-4. c. A speculative visual representation of the human VN cross-section showing the hypothesised location of cardiac efferent fibres (red), cardiac afferent fibres (blue), and laryngeal fibres (green).

The strongest activation of laryngeal EP was observed at channel 1 (Fig. 7b), reaching 63 µV, whereas the weakest activation occurred at channel 12, reachingan amplitudeof 31 µV. EP distribution suggests a higher density or number of laryngeal-specific fibres in the proximity of channels 1 to 4. Physiological effects of laryngeal and cardiac stimulation show fibre separation of 231° around the circumference of the VN (Fig. 7c).

## 4. Discussion

Design, manufacturing, testing and validation, animal test, and a first pilot human trial are reported for the 15-channel selective VN stimulator. The stimulator was designed with safety and channel selectivity in mind, thus achieving sVNS with full 15-channel selective cuff compatibility. The proposed system has a battery-free, fully wireless architecture with a multichannel output stage capable of selectively addressing individual stimulation channels. NFC is used both for communication and to power the device through NFC power harvesting. The device reliably generates stimulation pulses according to the programmed parameters. The worst-case percentage error was 3.83% for current amplitude, 10.88% for PW, and 6.04% for pulse frequency.

The stimulator was tested on a bench and verified in an animal model *in vivo*, where it achieved a selective bradycardia of 23.28 ± 12.91% when stimulating efferent porcine cardiac fibres.

The study demonstrates the feasibility of sVNS using an implantable sVNS system to modulate the activity of the PNS. This is validated through in vivo porcine experiments and a first-in-human pilot application. The manufactured sVNS device achieved consistency, reproducibility, and a similar level of fascicle selectivity (Fig. 5b and supplemental materials Fig. S2 and S3) to the previously reported results obtained using a benchtop system [23]. This highlights function-specific fibre distribution in pigs, allowing the possibility of targeting specific physiological function by tapping into neural pathways from the cervical section of the VN.

Similarly, a pilot application of sVNS in the human demonstrated the feasibility of eliciting a clinically relevant biomarker response (HR change) separately and in a different location from a negative side effect-associated one (largest laryngeal EP) (Fig. 7a and 7b). While different activation distributions are observed for HR (Fig. 7a) and laryngeal EP (Fig. 7b), it remains unclear whether the observed reduction in laryngeal activation is sufficient to meaningfully reduce the perception of stimulation-induced side effects. Furthermore, we do not define explicit selectivity thresholds for either laryngeal EP or HR reduction, as the degree of HR reduction required for a therapeutic effect in heart failure and the threshold at which reductions in laryngeal activation translate into clinically meaningful improvements in side-effect perception have yet to be established.

A clear separation between bradycardic and tachycardic effects was observed (Fig. 7a). Previous studies in pigs suggest that stimulation of the efferent fibres leads to a decrease in HR, whereas stimulation of afferent fibres increases HR [1]. Fig. 7a shows the potential distinction in these fibres: stimulation of channels 1 to 7 and 11 to 14 shows a mild increase in HR, aligning with the cardiac afferent fibres, whereas stimulation of channels 8 to 10 results in a prominent decrease of HR, with the highest decrease of 7.5%. The potential recruitment of cardiac efferent fibres suggests a possible therapeutic target for heart failure, addressing a key limitation of conventional VNS approaches, including the NECTAR trial, which failed to demonstrate prevention of cardiac remodelling [21]. Furthermore, peaks in HR modulation and laryngeal evoked activity were observed in diametrically opposite channels, indicating the potential to minimise laryngeal side effects while preserving the ability to recruit cardiac fibres. Activation of putative cardiac afferent fibres, which leads to an increase in HR in some channels, was observed in the human experiment and was not observed in the porcine trial. One of the factors contributing to this effect could be the apparent difference in the anaesthetic regimen. A combination of tiletamine-zolazepam, fentanyl, and isoflurane was administered to the pigs, whereas propofol and sevoflurane were used for induction and maintenance of anaesthesia in the human trial. The choice of anaesthetic agents in the porcine trial was specifically dictated by the lack of vagosuppressive and parasympathetic effects, whereas the anaesthetic regimen in the human trial was determined by the clinical requirements of the surgical procedure and could not be modified for this study.

On the other hand, while the porcine model is considered a good model of the human VN, recent anatomical studies have shown that the difference in fascicular organisation can be substantial. Variability in the human VN structure between subjects also seems to be large [24,38]. The autonomic reflex pathways, such as the baroreflex, Bainbridge reflex, and Hering-Breuer reflex, may be differentially recruited in humans and pigs, resulting in species-specific predominance of bradycardic versus tachycardic responses during in vivo experiments. Direct evidence comparing the relative activation of these reflexes between species remains limited, and further research is needed to clarify their respective roles. While these findings remain preliminary, they demonstrate the potential of selective VNS to improve the development of VNS-based therapies for heart failure and may represent an important step in the transition from non-selective to function-specific stimulation approaches.

The device performs comparably to other implantable stimulators (Table 5), achieving the high channel count and voltage compliance essential for sVNS. For instance, this work shares the capability of wireless power and communication through the NFC interface with the ReStore device due to a similar power, MCU, and communication stage, while providing selectivity in the output stage suitable for the application [31]. Comparison with the benchtop ScouseTom system used in the previous sVNS study demonstrates comparable reductions in HR [27]. Similar HR percentage change distributions were observed (Supplementary Fig. S4). Statistical analysis of the porcine trial did not reveal a statistically significant difference between the benchtop and the implantable system when using the same multielectrode array type in the same pig cohort (Supplementary Table S1).

**Table 5.** Comparison with other relevant works.

| Device (Ref.) | Application | Power Source | Package Volume (mm<sup>3</sup>) | Channels | PW Range (μs) | Maximum Current (mA) | Voltage Compliance (V) |
| --- | --- | --- | --- | --- | --- | --- | --- |
| OpenNerve [48] | IPG<sup>1</sup>, recorder, biomarker sensor | 2× rechargeable batteries | 31,123.6 | 4×4-contact or 2×8-contact connectors | 100-1000 | 5 | ±8 |
| Implantable bioelectronics for gut electrophysiology [49] | Recording (MEA<sup>2</sup>) | Intan Headstage (percutaneous) | Not directly comparable* | 7 (tetrodes) | N/A | N/A | N/A |
| Ultrasonically Powered Neuromodulation Platform [50] | PNS<sup>3</sup>/VNS<sup>4</sup>/sVNS<sup>5</sup> | Wireless ultrasonic | Not reported (benchtop) | 2 inhibition, 6 excitation | 0-1500 | Voltage-controlled | ±4V |
| Livanova VNS Therapy System | VNS | Non-rechargeable battery | 9,936 (+ lead) | 1 | 130-1000 | 3.5 | Not reported |
| Magnetoelectric implant [51] | PNS | Wireless magnetoelectric | 95.46 | 1 | 50-1200 | Voltage-controlled | 0.3-3.3 |
| ReStore [31] | PNS | NFC<sup>6</sup> power harvesting | 312 | 1 | 100-250 | 1.2 | 12 |
| Versatile Hermetic Implant [40] | PNS | Wireless power harvesting | 5,687<sup>7</sup> | 36 | 0-500 | 3 | 16 |
| This work | sVNS | NFC power harvesting | 9,690.5 | 15 | 50-4000 | 2.026 | ±20 |

<sup>1</sup>IPG – Implantable Pulse Generator<sup>2</sup>MEA – Multielectrode Array<sup>3</sup>PNS – Peripheral Nervous System<sup>4</sup>VNS – Vagus Nerve Stimulation<sup>5</sup>sVNS – selective Vagus Nerve stimulation<sup>6</sup>NFC – Near-Field Communication protocol<sup>7</sup>Encapsulated in ceramic cover, excluding separate power and data coils
\*Recording-only system; volume not directly comparable to fully implantable stimulators

The accelerated ageing tests suggest that while the current device architecture is suitable for acute studies in both pigs and humans, it is susceptible to corrosion. The device would require substantial changes to the packaging strategy to remain functional on the scale of years to decades as a fully implantable device.

## 5. Limitations and future work

One limitation of this work is the reliance on a commercially available NFC transmitter device based on an Arduino shield. The current combination of the sVNS pulse generator and the transmitter yields a very modest communication and power range. The working range of 10 mm was sufficient for our tests, accelerating ageing setup, in vivo acute porcine trials, and first human trials with the modified clinical sVNS setup, but further wireless range extension will be necessary for chronic experiments with freely moving animals.

The receiver antenna of the sVNS pulse generator is designed using a tightly wrapped copper wire. This setup inherently introduces variability in coil inductance and capacitor reactance, and therefore lacks scalability. Future work will include improving device scalability through the development of a tunable PCB-trace antenna and transmitter capable of delivering the required power over longer distances *in vivo*.

Charge imbalance observed after benchtop biphasic injections is a current source characteristic, which we attributed to a combination of the operational amplifier slew rate (4.5 V/µs), gain-bandwidth product (1.1 MHz), settling time (up to 5 µs), switching circuit break-before-make delays, and DC blocking capacitor discharge cycles. The largest net injected charge, expressed as a percentage of the nominal biphasic pulse charge in 0.9% saline, was -1.86% and 3.13% for 50 µs at 1 mA and 1 ms at 1 mA, respectively.

The accelerated ageing setup would be better informed by a higher number of samples. Nevertheless, few published ageing studies demonstrate functional devices under stress. Studies typically age test structures meant to mimic parts of devices, such as interconnects, or insulation [39]. Future work would explore ways to make the packaging more robust, such as using high-reliability ceramic hybrid circuits and hermetic packaging with improved underwater adhesion to silicone rubber encapsulation, reducing the risk of device failure from corrosion [40,41].

A trial-and-error stimulation paradigm was used to identify the optimal electrode pair. An embedded on-chip imaging technique, such as fast neural electrical impedance tomography, could potentially localise the nerve fibres of interest and thereby avoid the need for a trial-and-error approach [42,43].

The human patient was anaesthetised using a combination of propofol and other drugs, which are chosen at the anaesthetist’s discretion. The choice and combination of anaesthetic drugs are of very limited control from the pilot research perspective. Propofol is a known anaesthetic agent that desensitises muscular responses and the activity of the PNS in general [44–46]. Further limitation of the pilot human trial is the length of the experiment. The trial was performed as a preamble to clinically approved Livanova SenTiva VNS surgery, which limited the experiment duration to 30 minutes.

Due to time constraints and variability in autonomic responses during the surgical procedure, the stimulation-induced bradycardic response was compared to the stimulation-free period of the respective channel. A more objective comparison could be achieved by contrasting the HR decrease with the average HR recorded while the patient was in a stable anaesthetised state. Further increase in the experiment time may be necessary to enable additional stimulation paradigms aimed at improving fascicle localisation and recruitment.

The system was employed in an n = 1 pilot human trial, which inherently lacked sufficient statistical power. A larger sample of patients is required to determine whether the observed stimulation effects are statistically significant.

Further porcine *in vivo* studies are planned to transition from the acute to a medium-term, chronically implanted model in freely moving animals, with the aim of evaluating heart failure outcome improvements.

## 6. Conclusion

We have demonstrated the design and testing of a precise and controllable miniature sVNS system, validated through benchtop, animal, and human tests. The sVNS hardware design files, firmware, and the GUI are openly available in a public repository [47]. Open-source availability of the hardware and the software enables the replication of the device and supports further sVNS studies. The sVNS system was shown to modulate cardiac activity in the porcine model and in the pilot human experiment, demonstrating translation from animal experiment to the human trial. Stimulation results provide evidence for spatially selective recruitment of cardiac fibres, allowing differentiation between bradycardic and tachycardic responses, while minimising the effects of laryngeal activation. This strategy can potentially be utilised to improve neuromodulation approaches for heart failure. In comparison with the existing stimulation technologies, the developed sVNS system provides a high output channel count, voltage compliance of ±20 V, which is sufficient for stimulation through platinised stainless-steel electrodes, wireless power and communication, a graphical user interface, and biocompatible silicone packaging suitable for short-to-medium-term implantation.

## Acknowledgements

This work was supported by the following grants: EPSRC EP/X018415/1, MRC MR/Z504555/1, NIH SPARC 3OT2OD026545, UCL spinout Cyqiq Ltd, and Kirill Aristovich.

Author E. Ravagli is currently supported by a Horizon MSCA Guarantee Postdoctoral Fellowship (UKRI Grant EP/X03691X/1). Olujimi Ajijola and Kalyanam Shivkumar were supported by Leducq Foundation grant on Bioelectronic medicine.

## Ethics statement

This study was performed in accordance with the Declaration of Helsinki. This human study was approved by NHS Research Ethics Committee - approval: 22/LO/0463. All adult participants provided written informed consent to participate in this study. This animal study was approved by UCLA Chancellor’s Animal Research Committee - approval: ARC 2016 085. The study was performed within the guidelines set by the University of California Institutional Animal Care and Use Committee (IUCUC) and the National Institutes of Health Guide for Care and Use of Laboratory Animals. Experiments were approved by the UCLA Chancellor’s Animal Research Committee. All authors understand the ethical principles under which the journal operates, and the work presented here complies with animal ethics.

