## Supplementary Figures and Tables for "An Implantable Wireless Battery-Free Spatially Selective Vagus Nerve Stimulator"

### Supplementary materials

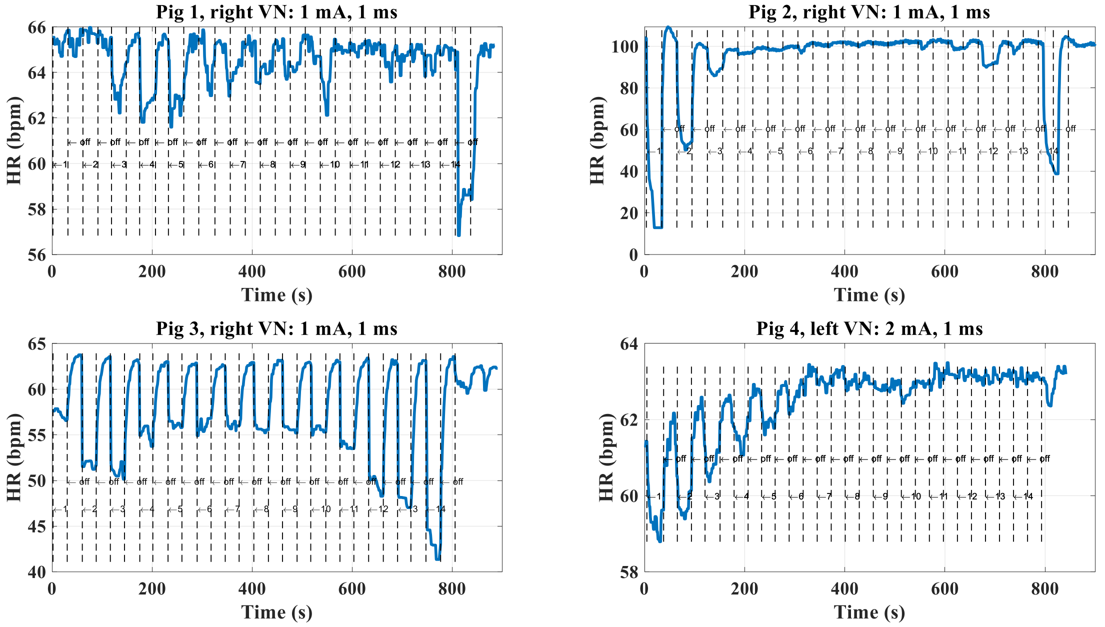


**Figure S1. Raw heart rate recorded during the acute porcine cardiac trial when utilising the implantable selective vagus nerve stimulation device. All stimulations were delivered at 20 Hz.**

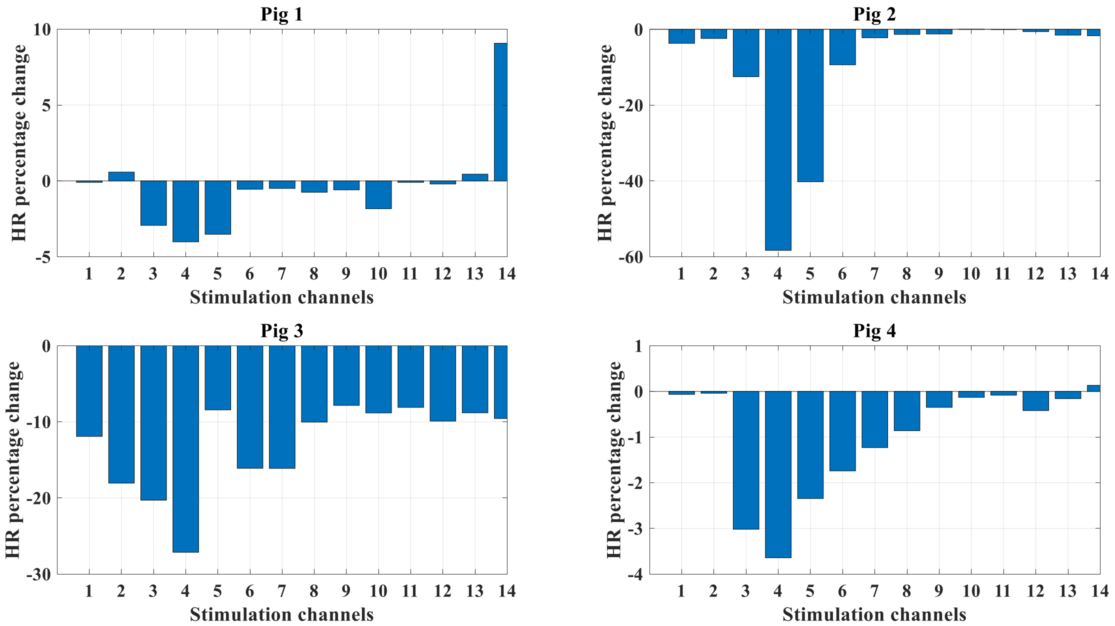


**Figure S2. Heart rate (HR) percentage change of the same pigs as in Fig. S1 after aligning the largest HR decrease with channel 4.**

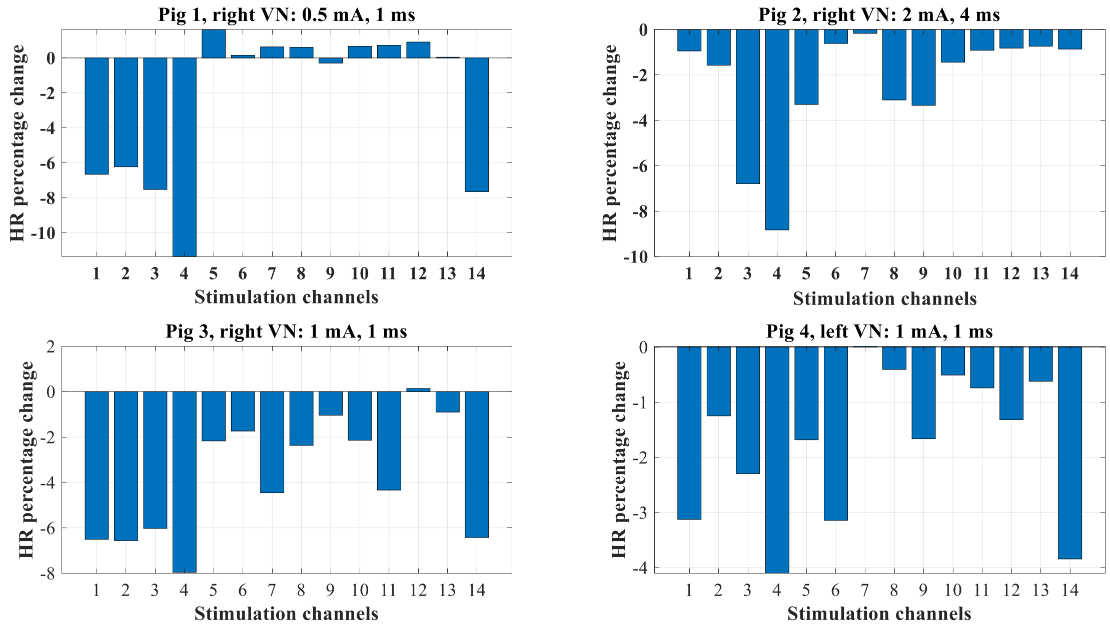


**Figure S3. Heart rate (HR) percentage change of the same pigs as in Fig. S1, elicited utilising a benchtop ScouseTom system, after aligning the largest HR decrease with channel 4.**

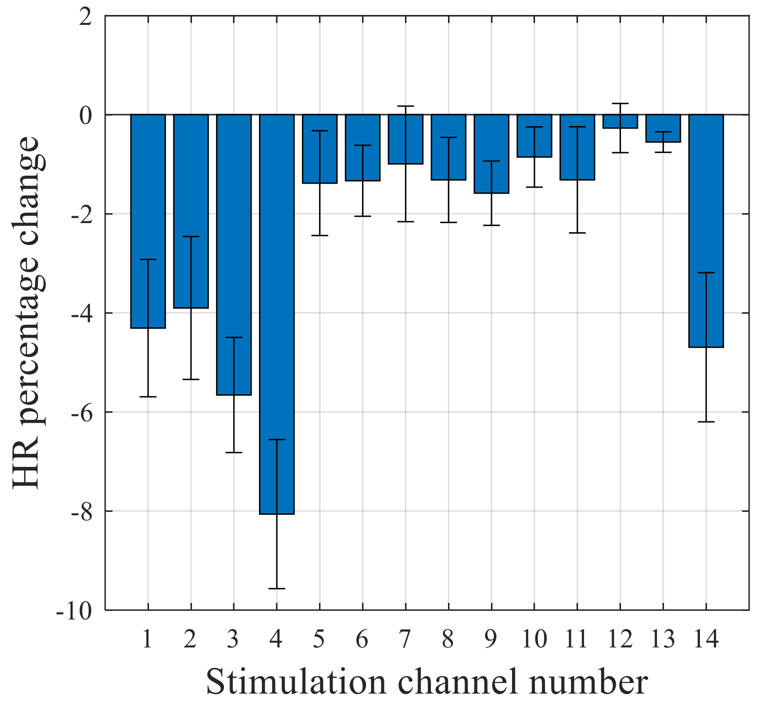


**Figure S4. Mean heart rate (HR) percentage change ± standard error of the mean, elicited utilising a benchtop ScouseTom system, across all porcine trials (n = 4), after aligning the maximal HR decrease with channel 4.**

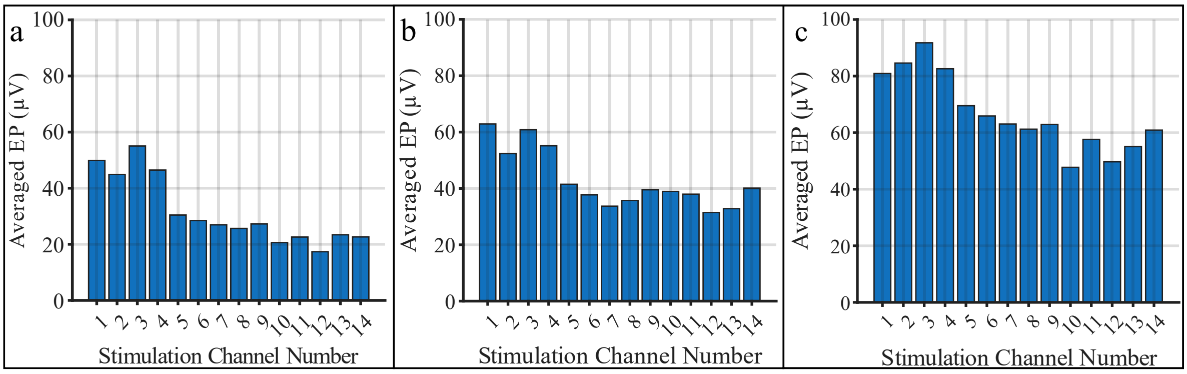


**Figure S5. Laryngeal motor evoked potentials calculated from different available EMG channels: a – forehead, b – chin, c – neck.**

**Table S1. Paired-sample t-test p-values comparing the mean heart rate change evoked by stimulation of corresponding channels using the benchtop ScouseTom system (Figure S4) and the implantable sVNS system (Figure 5b in the main text), across all porcine trials (n = 4).**

| Channel No | P-value |
|---|---|
| 1 | 0.9006 |
| 2 | 0.7978 |
| 3 | 0.3898 |
| 4 | 0.3179 |
| 5 | 0.2371 |
| 6 | 0.2222 |
| 7 | 0.2138 |
| 8 | 0.4122 |
| 9 | 0.6834 |
| 10 | 0.3920 |
| 11 | 0.5339 |
| 12 | 0.3984 |
| 13 | 0.4017 |
| 14 | 0.4145 |
